# Cefepime-Taniborbactam and Ceftibuten-Ledaborbactam Maintain Activity Against KPC Variants that Lead to Ceftazidime-Avibactam Resistance

**DOI:** 10.1101/2024.10.11.617870

**Authors:** Cullen L. Myers, Annie Stevenson, Brittany Miller, Denis M. Daigle, Tsuyoshi Uehara, Daniel C. Pevear

**Affiliations:** Department of Molecular and Cellular Biology, University of Guelph, Guelph, Ontario, Canada; Venatorx Pharmaceuticals Inc., Malvern, Pennsylvania, USA

**Author notes:** Address correspondence to:* Cullen L. Myers.

**Keywords:** taniborbactam, ledaborbactam, KPC, beta-lactamase inhibitor, ceftazidime-avibactam, antibiotic resistance

## Abstract

*Klebsiella pneumoniae* carbapenemases (KPCs) are widespread β-lactamases that are a major cause of clinical non-susceptibility of Gram-negative bacteria to carbapenems and other β-lactam antibiotics. Ceftazidime combined with the β-lactamase inhibitor avibactam (CAZ-AVI) has been effective for treating infections by KPC-producing bacteria, but emergent KPC variants confer resistance to the combination. Taniborbactam and ledaborbactam are bicyclic boronate β-lactamase inhibitors under development with cefepime and ceftibuten, respectively, to treat carbapenem-resistant bacterial infections. Here, we assessed the effects of clinically important KPC-2 and KPC-3 variants (V240G, D179Y, D179Y T243M) on the antibacterial activity of cefepime-taniborbactam (FEP-TAN) and ceftibuten-ledaborbactam (CTB-LED) and examined catalytic activity and inhibition of these variants. FEP-TAN and CTB-LED were highly active against CAZ-AVI-resistant engineered *E. coli* strains expressing these variants. Purified KPC variants catalyzed more efficient CAZ hydrolysis than wild-type enzymes, and D179Y-containing KPC-3 variants additionally catalyzed more efficient FEP hydrolysis than wild-type KPC-3. All KPC variants poorly hydrolyzed CTB, and D179Y-containing variants demonstrated significantly higher affinity for CAZ than FEP or CTB. Second-order rate constants (*k*^2^/*K*) for inhibition of D179Y-containing KPC-2 variants were significantly reduced relative to wild-type KPC-2, with AVI most impacted. *K*^2^/*K* was less affected for D179Y-containing KPC-3 variants, and reflected robust inhibition by TAN, LED and AVI. Together, the findings illustrate a biochemical basis for greater FEP-TAN and CTB-LED antibacterial activity in KPC variant expression backgrounds relative to CAZ-AVI, whereby the boronate inhibitors have sufficient inhibitory activity, whilst FEP and CTB are poorer substrates and bind to the variant enzymes with reduced affinity compared to CAZ.

## Introduction

β-lactam (BL) antibiotics continue to be the first-line therapy for the treatment of Gram-negative bacterial infections due to their proven track record of clinical safety and efficacy^1^. Among the various mechanisms of bacterial resistance to these drugs, the production of β-lactamase enzymes that hydrolyze the β-lactam ring is foremost^1^. Of particular concern are carbapenemases, including globally disseminating *Klebsiella pneumoniae* carbapenemases (KPCs) that inactivate so-called “last line” carbapenem antibiotics (e.g., meropenem). Carbapenemases are distributed throughout the four Ambler Classes – classes A, C and D utilize a nucleophilic serine to catalyze BL hydrolysis (serine β-lactamases), while class B enzymes (metallo β-lactamases) rely on active site zinc ions^2^. KPCs belong to Class A, and hydrolysis initiates with the attack of the BL ring by the catalytic serine, forming an acyl-enzyme complex that is subsequently deacylated to release an inactive product^3^. The substrate profile for KPCs encompasses virtually all β-lactams antibiotics, which limits therapeutic options for infections by KPC-producing bacteria rendering such infections a serious global health threat^4,5^.

To address β-lactamase-mediated antibiotic resistance, β-lactamase inhibitors (BLIs) have been deployed to the clinic, with demonstrable success at rescuing BL efficacy in treating infections by β-lactamase producing bacteria^6^. KPCs, however, are refractory to early-generation BLIs that possess the BL moiety, e.g., clavulanic acid and tazobactam^7,8^. This spurred the development of BLIs derived from non-BL scaffolds that achieve clinically relevant inhibition of KPCs^9,10^. The diazabicyclooctane (DBO) avibactam (AVI) was the first such BLI to enter the clinic for treatment of carbapenem-resistant *Enterobacterales* infections. The combination product ceftazidime-avibactam (Avycaz^®^) received FDA approval in 2015^11,12^ and in 2017, the first (mono)cyclic boronate BLI, vaborbactam, was approved for clinical use in combination with meropenem (Vabomere^®^)^13^.

Avycaz^®^ was a welcome introduction to the clinic that restored CAZ efficacy against KPC-producing bacteria with the inhibitor avibactam. However, concurrent with and following its approval for clinical use, the combination was shown to select for CAZ-AVI resistance linked to *bla*^KPC^ mutations, *in vitro*^14,15^. The Ω-loop region of the protein, which spans residues 162-179 in KPC, has proved to be a hotspot for amino acid changes resulting from such mutations in *bla*^KPC^, a prime example being substitutions at D179^14,16,17^. Mutations causing amino acid substitutions to residues outside the Ω-loop have also been associated with elevated CAZ-AVI MICs. For instance, V240 is in a region adjacent to the Ω-loop and changes at this site have been proposed to affect Ω-loop dynamics^18^. Worryingly, KPC variants associated with CAZ-AVI resistance have since emerged during the course of therapy with Avycaz^®19,20^.

The Ω-loop is a conserved feature among Class A β-lactamases that is proximal to the KPC active site, with loop residues E166 and N170 oriented to position an active site water molecule critical for the deacylation step of substrate hydrolysis^21^. Mutations that alter residues within the KPC Ω-loop have varied impacts on the antibacterial activity of BLs or BL-BLI combinations^22–24^. Structural and biochemical evidence suggests that certain amino acid substitutions at D179 cause increased CAZ affinity for the enzyme, evoking a trapping mechanism in which such variants sequester the antibiotic away from binding to its cellular penicillin-binding protein targets^22,24^. Such variants poorly hydrolyze carbapenems, and strains harboring these KPC variants are susceptible to meropenem^19,20,25^. This complicates therapeutic strategies since infections initially present as MEM-resistant but revert to MEM-susceptible while becoming CAZ-AVI-resistant following treatment with the combination^19^. Substitutions at D179 also reportedly directly affect the inhibitory activity of AVI and VAB, the latter being less affected^23,24,26^.

Taniborbactam is a bicyclic boronate BLI being developed with cefepime (FEP) for the treatment of complicated urinary tract infections, as well as hospital- and ventilator-associated bacterial pneumonia infections caused by β-lactamase-producing Enterobacterales and *Pseudomonas aeruginosa*^27–29^. Ledaborbactam (LED) is the active form of the orally bioavailable bicyclic boronate prodrug ledaborbactam-etzadroxil that is in development with ceftibuten (CTB) to address multi-drug-resistant β-lactamase-expressing Enterobacterales^30,31^. TAN and LED both provide excellent coverage of class A (including KPCs) and class C serine β-lactamases, whereas the inhibition spectrum for TAN also includes clinically relevant variants of the metallo β-lactamases NDM and VIM^28^.

In this study, we compared the antibacterial activity of FEP-TAN, CTB-LED and CAZ-AVI in isogenic *E. coli* strains producing KPC-2 and KPC-3 variants implicated in CAZ-AVI resistance and examined the catalytic properties and inhibition of these variants.

## Results

### Effect of KPC-variant expression on antibacterial activity

Minimum inhibitory concentrations (MICs) for BLs and BL-BLI combinations were determined against isogenic *E. coli* strains expressing KPC-2 or KPC-3 variants (V240G, D179Y and D179Y T243M; Fig. 1) known to confer CAZ-AVI non-susceptibility (Tables 1 and 2). KPC-2^V240G^ expression increased the MICs for BLs relative to the control strain lacking KPC expression, and TAN, AVI or LED fully restored FEP or CTB activity, while CAZ-AVI and CAZ-TAN MICs remained elevated at 4 – 8 µg/mL (Table 1). Owing to previously described collateral sensitivity towards carbapenems^15,19^, MICs for MEM were modestly increased (8-fold) by the expression of KPC-2^D179Y^ or KPC-2^D179Y^ ^T243M^ relative to the control strain, compared to increases of > 512-fold for CAZ, 64-fold for FEP and 8-16-fold for CTB. MICs for CAZ-AVI remained highly elevated against these strains (> 256 µg/mL), but FEP-TAN and CTB-LED were active at ≤ 0.5 µg/mL. FEP-AVI was also active against the strains expressing KPC-2^D179Y^ or KPC-2^D179Y^ ^T243M^ at MICs of 2 µg/mL and 0.5 µg/mL, respectively (Table 1), possibly reflecting the greater FEP stability to the KPC variants, compared to CAZ. Indeed, FEP was active at 16 µg/mL against these stains, compared to MICs of > 256 µg/ mL for CAZ. Moreover, TAN only rescued CAZ MICs to 16 µg/ml against *E. coli* producing KPC-2^D179Y^ or KPC-2^D179Y^ ^T243M^ (Table 1).

**Figure 1.**
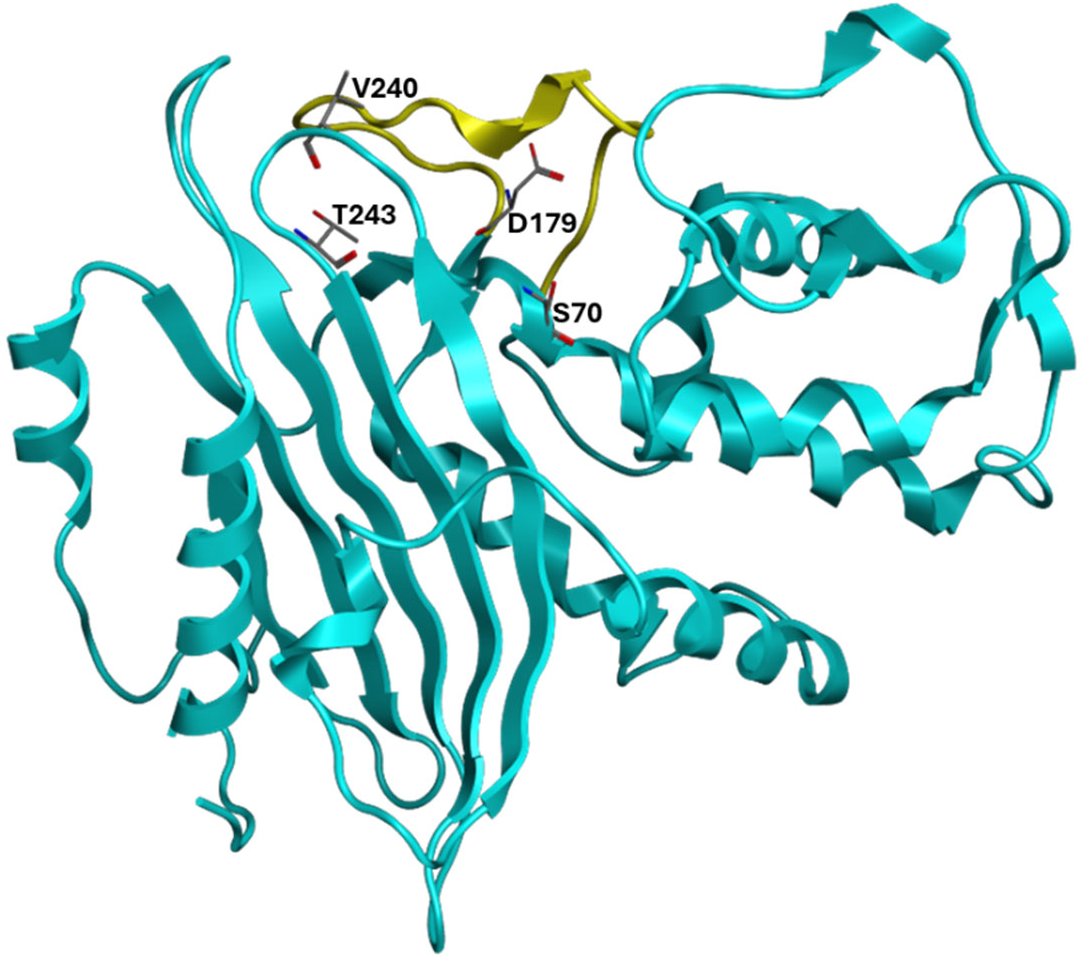
Amino acid residues implicated in KPC variant-mediated CAZ-AVI resistance. The X-ray crystal structure of KPC-2 (PDB: 5UL8^32^) is shown. The catalytic serine (S70), and residues altered by *bla*^KPC^ mutations associated with clinical CAZ-AVI resistance are shown as sticks. The Ω-loop is rendered in yellow.

**Table 1.** Antibacterial activity (MIC, μg/mL) of BLs and BL/ BLIi^a^ combinations against engineered *E. coli* strains expressing wild-type KPC-2 or KPC-2 variants, relative to a control strain lacking a β-lactamase.

| KPC-2 variant expressed | MEM | CAZ | CAZ+ AVI | CAZ + TAN | FEP | FEP + TAN | FEP + AVI | CTB | CTB + LED |
| --- | --- | --- | --- | --- | --- | --- | --- | --- | --- |
| None | 0.03 | 0.5 | 0.5 | 0.5 | 0.12 | 0.12 | 0.12 | 1 | 0.25 |
| wt | 32 | 64 | 1 | 0.5 | 64 | 0.12 | 0.25 | 16 | 0.25 |
| V240G | 8 | > 256 | 8 | 4 | 128 | 0.25 | 0.25 | 64 | 0.5 |
| D179Y | 0.25 | > 256 | 64 | 16 | 16 | 0.5 | 2 | 16 | 0.5 |
| D179Y T243M | 0.25 | > 256 | 64 | 16 | 16 | 0.25 | 0.5 | 16 | 0.25 |

<sup>a</sup>BLs were tested with BLIs at a fixed concentration of 4 µg/mL.

**Table 2.** Antibacterial activity (MIC, μg/mL) of BLs and BL/ BLIi^a^ combinations against engineered *E. coli* strains expressing wild-type KPC-3 or KPC-3 variants, relative to a control strain lacking a β-lactamase.

| KPC-3 variant expressed | MEM | CAZ | CAZ + AVI | CAZ + TAN | FEP | FEP + TAN | FEP + AVI | CTB | CTB + LED |
| --- | --- | --- | --- | --- | --- | --- | --- | --- | --- |
| None | 0.03 | 0.5 | 0.5 | 0.5 | 0.12 | 0.12 | 0.12 | 1 | 0.25 |
| wt | 16 | 256 | 4 | 2 | 128 | 0.25 | 0.5 | 32 | 0.5 |
| V240G | 16 | > 256 | 128 | 16 | 256 | 0.5 | 1 | 32 | 0.5 |
| D179Y | 0.12 | > 256 | 256 | 32 | 32 | 1 | 4 | 32 | 1 |
| D179Y T243M | 0.12 | > 256 | 256 | 32 | 16 | 0.5 | 2 | 16 | 0.5 |

<sup>a</sup>BLs were tested with BLIs at a fixed concentration of 4 µg/mL

The MIC for CAZ-AVI was elevated 256-fold against the KPC-3^V240G^-expressing strain relative to the control strain, whereas TAN and LED fully restored FEP and CTB activity, respectively (Table 2). TAN only partially rescued CAZ activity in this strain (MIC of 16 µg/mL), but it is notable that the MIC of CAZ was highly elevated in the absence of a BLI for this strain (> 256 µg/mL). FEP activity, however, was rescued to a MIC of 1 µg/mL by AVI (128 µg/mL in the absence of BLI).

Like the D179Y-containing KPC-2 variants, strains expressing D179Y-containing KPC-3 variants were susceptible to MEM, and MICs for CAZ-AVI were highly elevated at 256 µg/mL, while FEP-TAN and CTB-LED were active at MICs of 0.5 – 1 µg/mL. AVI achieved appreciable rescue of FEP activity against these strains with MICs of 0.5 – 4 µg/mL, but CAZ-TAN activity was markedly weaker at 16 µg/mL (Table 2). Again, CAZ MICs in the absence of a BLI were significantly higher than FEP or CTB for these strains.

Overall, these results confirm our previous reporting of potent antibacterial activity for FEP-TAN or CTB-LED against CAZ-AVI-resistant KPC variant-producing strains^28,31^, while alluding to different effects of the substitutions on the hydrolytic activity and inhibition of KPC-2 versus KPC-3.

### Catalytic activity of KPC variants

Prompted by the differential effects of KPC-2/ -3 variant production on antibacterial activity, we determined the kinetic parameters for BL substrate hydrolysis using purified KPC enzymes. Kinetic parameters for hydrolysis of all substrates tested with KPC-2^V240G^ were comparable to KPC-2^wt^ (Table 3). In contrast to linear kinetics for CAZ hydrolysis by KPC-2^wt^ and KPC-2^V240G^, the D179Y-containing KPC-2 variants displayed saturation kinetics for CAZ hydrolysis, with reduced *K*^M^ and increased *k*^cat^/ *K*^M^ relative to KPC-2^wt^ (Table 3) consistent with previous reports^23,26^. *K*^M^ for FEP hydrolysis was also reduced with these variants, but unlike CAZ, catalytic efficiency (*k*^cat^/*K*^M^) decreased relative to wild-type KPC-2, and both variants poorly hydrolyzed CTB (Table 3). Interestingly, KPC-2^D179Y^ or KPC-2^D179Y^ ^T243M^ hydrolyzed FEP with a higher *k*^cat^ /*K*^M^ than CAZ, yet FEP had lower MICs against strains expressing these variants (Table 1, Table 3). Still, the findings are in agreement with reports of elevated CAZ MICs against strains expressing KPC variants with D179 substitutions that appeared to have reduced hydrolytic activity against this substrate^23,24,26^.

**Table 3.** Kinetic parameters for β-lactam hydrolysis by KPC-2 variants.

| Parameter | WT | V240G | D179Y | D179Y T243M |
| --- | --- | --- | --- | --- |
| **Nitrocefin** |  |  |  |  |
| $K_M$ ($\mu\text{M}$) | $15.2 \pm 1.2$ | $14.3 \pm 0.2$ | $5.0 \pm 2.1$ | $2.3 \pm 0.75$ |
| $k_{\text{cat}}$ ($\text{sec}^{-1}$) | $38.8 \pm 2.2$ | $38.5 \pm 0.6$ | $0.13 \pm 0.02$ | $(0.61 \pm 0.05) \times 10^{-2}$ |
| $k_{\text{cat}}/K_M$ ($\text{sec}^{-1}.\text{M}^{-1}$) | $(2.4 \pm 0.14) \times 10^6$ | $(2.4 \pm 0.4) \times 10^6$ | $(8.2 \pm 1.4) \times 10^3$ | $(3.8 \pm 0.32) \times 10^2$ |
| **CENTA<sup>33</sup>** |  |  |  |  |
| $K_M$ ($\mu\text{M}$) | $19.7 \pm 1.3$ | $63.3 \pm 2.3$ | $1.9 \pm 0.2$ | $2.1 \pm 0.3$ |
| $k_{\text{cat}}$ ($\text{sec}^{-1}$) | $243 \pm 11$ | $662 \pm 9.6$ | $(1.2 \pm 0.08) \times 10^{-2}$ | $(0.9 \pm 0.04) \times 10^{-2}$ |
| $k_{\text{cat}}/K_M$ ($\text{sec}^{-1}.\text{M}^{-1}$) | $(1.5 \pm 0.07) \times 10^7$ | $(4.1 \pm 0.06) \times 10^7$ | $(7.4 \pm 0.5) \times 10^2$ | $(5.5 \pm 0.3) \times 10^5$ |
| **Ceftazidime** |  |  |  |  |
| $K_M$ ($\mu\text{M}$) | > 500 | > 500 | $0.6 \pm 0.09$ | $1.0 \pm 0.2$ |
| $k_{\text{cat}}$ ($\text{sec}^{-1}$) | - | - | $(1.2 \pm 0.04) \times 10^{-3}$ | $(3.7 \pm 0.13) \times 10^{-3}$ |
| $k_{\text{cat}}/K_M$ ($\text{sec}^{-1}.\text{M}^{-1}$) | $^a(7.3 \pm 1.3) \times 10^2$ | $^a(8.0 \pm 0.33) \times 10^2$ | $2.0 \times 10^3$ | $3.7 \times 10^3$ |
| **Cefepime** |  |  |  |  |
| $K_M$ ($\mu\text{M}$) | > 500 | > 500 | $3.7 \pm 0.7$ | $3.0 \pm 0.6$ |
| $k_{\text{cat}}$ ($\text{sec}^{-1}$) | - | - | $(1.5 \pm 0.07) \times 10^{-2}$ | $(1.7 \pm 0.06) \times 10^{-2}$ |
| $k_{\text{cat}}/K_M$ ($\text{sec}^{-1}.\text{M}^{-1}$) | $^a(1.3 \pm 0.04) \times 10^5$ | $^a(1.9 \pm 0.3) \times 10^5$ | $4.0 \times 10^3$ | $5.7 \times 10^3$ |
| **Ceftibuten** |  |  |  |  |
| $K_M$ ($\mu\text{M}$) | > 500 | > 500 | ND | ND |
| $k_{\text{cat}}$ ($\text{sec}^{-1}$) | - | - | ND | ND |
| $k_{\text{cat}}/K_M$ ($\text{sec}^{-1}.\text{M}^{-1}$) | $^a(2.2 \pm 0.24) \times 10^3$ | $^a(2.0 \pm 0.25) \times 10^3$ | ND | ND |

Data are reported as the mean ± standard deviation, (n = 3)
a: $k_{\text{cat}}/K_M$ was determined from plots of velocity versus substrate concentration, where $[S] \ll K_M$ <sup>34</sup>
ND, not determined as hydrolysis was below the detection limit.

Catalytic profiles for the KPC-3 variants were generally similar to those for the KPC-2 variants, with noteworthy differences (Table 4). The *K*^M^s for CENTA or nitrocefin hydrolysis by KPC-3^D179Y^ and KPC-3^D179Y/T243M^ were comparable to KPC-3^wt^, but *k*^cat^ was significantly reduced. Additionally, D179Y-containing KPC-3 variants displayed saturation kinetics for CAZ and FEP hydrolysis with reduced *K*^M^s but increased *k*^cat^/ *K*^M^ relative to KPC-3^wt^. Indeed, *K*^M^ and *k*^cat^ for CAZ hydrolysis by these variants were markedly lower than for FEP hydrolysis.

**Table 4.**
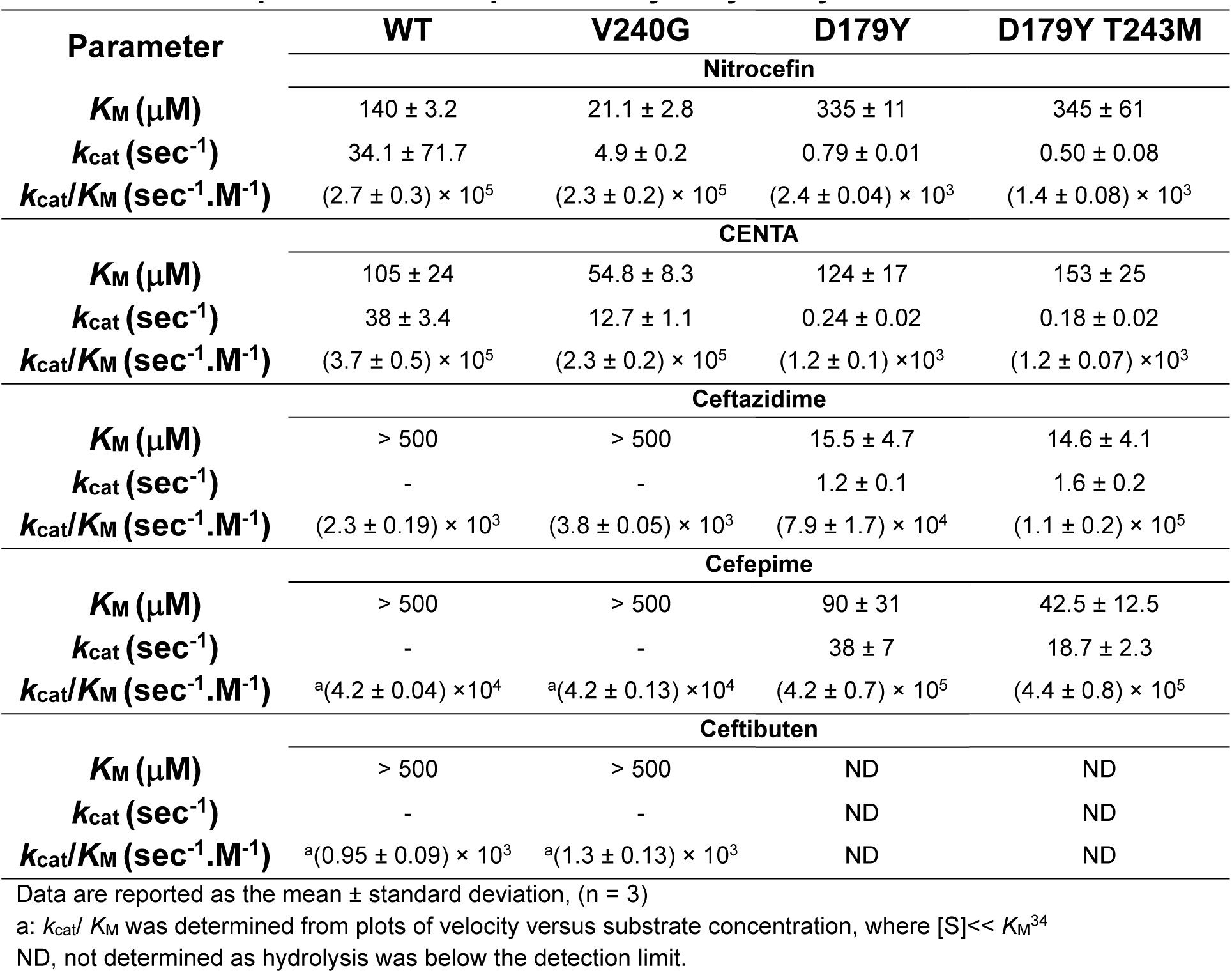
Kinetic parameters for β-lactam hydrolysis by KPC-3 variants.

| Parameter | WT | V240G | D179Y | D179Y T243M |
| --- | --- | --- | --- | --- |
| **Nitrocefin** |  |  |  |  |
| $K_M$ ($\mu\text{M}$) | 140 ± 3.2 | 21.1 ± 2.8 | 335 ± 11 | 345 ± 61 |
| $k_{\text{cat}}$ ($\text{sec}^{-1}$) | 34.1 ± 71.7 | 4.9 ± 0.2 | 0.79 ± 0.01 | 0.50 ± 0.08 |
| $k_{\text{cat}}/K_M$ ($\text{sec}^{-1}.\text{M}^{-1}$) | (2.7 ± 0.3) $\times 10^5$ | (2.3 ± 0.2) $\times 10^5$ | (2.4 ± 0.04) $\times 10^3$ | (1.4 ± 0.08) $\times 10^3$ |
| **CENTA** |  |  |  |  |
| $K_M$ ($\mu\text{M}$) | 105 ± 24 | 54.8 ± 8.3 | 124 ± 17 | 153 ± 25 |
| $k_{\text{cat}}$ ($\text{sec}^{-1}$) | 38 ± 3.4 | 12.7 ± 1.1 | 0.24 ± 0.02 | 0.18 ± 0.02 |
| $k_{\text{cat}}/K_M$ ($\text{sec}^{-1}.\text{M}^{-1}$) | (3.7 ± 0.5) $\times 10^5$ | (2.3 ± 0.2) $\times 10^5$ | (1.2 ± 0.1) $\times 10^3$ | (1.2 ± 0.07) $\times 10^3$ |
| **Ceftazidime** |  |  |  |  |
| $K_M$ ($\mu\text{M}$) | > 500 | > 500 | 15.5 ± 4.7 | 14.6 ± 4.1 |
| $k_{\text{cat}}$ ($\text{sec}^{-1}$) | - | - | 1.2 ± 0.1 | 1.6 ± 0.2 |
| $k_{\text{cat}}/K_M$ ($\text{sec}^{-1}.\text{M}^{-1}$) | (2.3 ± 0.19) $\times 10^3$ | (3.8 ± 0.05) $\times 10^3$ | (7.9 ± 1.7) $\times 10^4$ | (1.1 ± 0.2) $\times 10^5$ |
| **Cefepime** |  |  |  |  |
| $K_M$ ($\mu\text{M}$) | > 500 | > 500 | 90 ± 31 | 42.5 ± 12.5 |
| $k_{\text{cat}}$ ($\text{sec}^{-1}$) | - | - | 38 ± 7 | 18.7 ± 2.3 |
| $k_{\text{cat}}/K_M$ ($\text{sec}^{-1}.\text{M}^{-1}$) | <sup>a</sup>(4.2 ± 0.04) $\times 10^4$ | <sup>a</sup>(4.2 ± 0.13) $\times 10^4$ | (4.2 ± 0.7) $\times 10^5$ | (4.4 ± 0.8) $\times 10^5$ |
| **Ceftibuten** |  |  |  |  |
| $K_M$ ($\mu\text{M}$) | > 500 | > 500 | ND | ND |
| $k_{\text{cat}}$ ($\text{sec}^{-1}$) | - | - | ND | ND |
| $k_{\text{cat}}/K_M$ ($\text{sec}^{-1}.\text{M}^{-1}$) | <sup>a</sup>(0.95 ± 0.09) $\times 10^3$ | <sup>a</sup>(1.3 ± 0.13) $\times 10^3$ | ND | ND |

Data are reported as the mean ± standard deviation, (n = 3)
a: $k_{\text{cat}}/K_M$ was determined from plots of velocity versus substrate concentration, where $[S] \ll K_M$ <sup>34</sup>
ND, not determined as hydrolysis was below the detection limit.

Even though CTB MICs against strains expressing D179Y-containing KPC variants were elevated 16-32-fold compared to control strains (Tables 1 & 2), kinetic parameters for CTB hydrolysis could not be determined under the assay conditions. Progress curves confirmed these variants had hydrolytic activity with CTB, however turnover was significantly slower than with the wild-type enzymes (Figure 2). For the wild-type enzymes, hydrolysis neared completion within five minutes. By contrast, CTB hydrolysis by KPC-2 D179Y-containing variants started to approach completion at ∼ 50 minutes, while D179Y-containing KPC-3 variants achieved 30 – 40% CTB conversion within the same timeframe (Figure 2).

**Figure 2.**
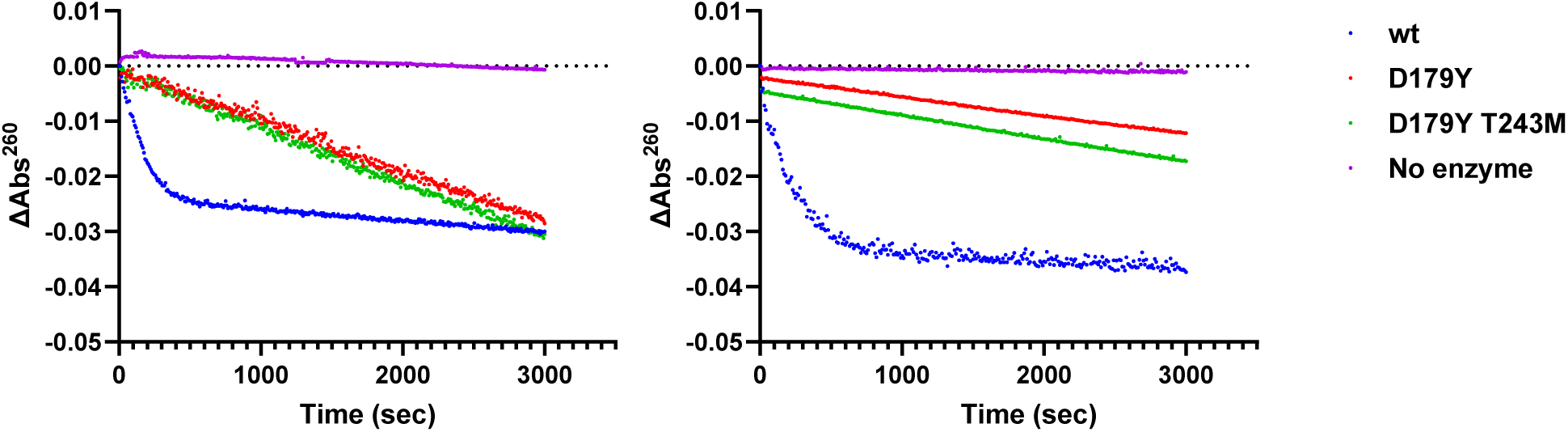
Hydrolysis of CTB by KPC variants. Progress curves are shown for reactions that contained 20 µM CTB and 1 µM of KPC-2 (left) or KPC-3 (right) variants.

A reduction in substrate turnover coupled with an apparent increase in affinity (i.e. reduced *K*^M^), as observed for CAZ hydrolysis by D179Y-containing variants, may indicate slow product release relative to the formation of enzyme-substrate complexes. The prolonged existence of an enzyme-substrate complex has been demonstrated for CAZ with the D179N variant of KPC-2^24^, and supported by pre-steady-state kinetic analyses of CAZ hydrolysis by KPC-2^D179Y^ ^35^. To assess whether the other cephalosporins used in this study possess higher affinities for D179Y-containing KPC variants than the wild-type enzymes, apparent *K*^i^ values for inhibition of CENTA hydrolysis by CAZ, FEP and CTB were determined (Table 5).

**Table 5.** Apparent affinities (*K*_i_^app^, µM) of CAZ, FEP and CTB for KPC variants.

|  | KPC Variant | Ceftazidime | Cefepime | Ceftibuten |
| --- | --- | --- | --- | --- |
| KPC-2 | wt | 19.8 ± 3.0 | 32.9 ± 5.6 | 159 ± 10 |
| | D179Y | 0.55 ± 0.12 | 4.6 ± 1.2 | ND |
| | D179Y/ T243M | 0.60 ± 0.19 | 3.2 ± 0.6 | ND |
| KPC-3 | wt | 437 ± 184 | 276 ± 47 | 368 ± 26 |
| | D179Y | 90 ± 3.1 | 1041 ± 72 | ND |
| | D179Y T243M | 177 ± 27 | 728 ± 58 | ND |

Data are reported as the mean ± standard deviation, (n = 3)
ND, no significant inhibition of CENTA hydrolysis was detected at the highest concentration tested.

CAZ and FEP showed significantly higher affinity for D179Y-containing KPC-2 variants relative to the wild-type enzyme, but FEP exhibited ∼ 10-fold weaker affinity for the variants than CAZ (Table 5). The apparent affinity of FEP for KPC-3^wt^ was comparable to that of CAZ, but FEP demonstrated weaker affinity for the D179Y-containing KPC-3 variants relative to KPC-3^wt^ (Table 5). Notably, CTB demonstrated weak affinity for wild-type KPC enzymes and lacked measurable inhibition of CENTA hydrolysis by KPC-2/ KPC-3 D179Y-containing variants.

### Inhibition of KPC variants

It was clear from data presented in Table 5 that the hydrolytic properties and BL affinities for KPC variants strongly influenced antibacterial activity BL-BLI combinations. However, the microbiological data could also be interpreted as indicating differing abilities of the BLIs to inhibit the KPC variants. For instance, TAN and LED rescued partner cephalosporins within 1 – 2 dilutions of the MICs for control strains, whereas MICs in the presence of AVI were invariably higher (see Tables 1 and 2). Therefore, to delineate the contributions from KPC variant inhibition to the observed antimicrobial susceptibilities, kinetic constants for inhibition of the variants by AVI, TAN and LED were determined.

AVI inactivated KPC V240G variants with reduced efficiencies compared to the wild-type enzyme (Tables 6 and 7). Dissociation rates were not impacted, and thus AVI *K*^d^ was increased for these variants. Similar observations were noted for TAN inhibition of KPC-2^V240G^ (Table 7). The *K*^d^ of TAN was also higher for KPC-3^V240G^ relative to the wild-type enzyme, which was in this case due to a faster dissociation rate since the second-order rate constant describing inactivation efficiency (*k*^2^/*K*) was not impacted (Table 7). These findings support the MIC results herein, along with previous findings showing that TAN and LED restored FEP or CTB antibacterial activity against *E. coli* expressing KPC-3^V240G28,36^.

**Table 6.** Kinetic parameters for inhibition of KPC-2 variants.

| Inhibitor | Parameter | wt | V240G | D179Y | D179YT243M |
| --- | --- | --- | --- | --- | --- |
| AVI | $k_2/K$ ($M^{-1}.sec^{-1}$) | $(3.6 \pm 0.2) \times 10^4$ | $(1.2 \pm 0.1) \times 10^4$ | $2.0 \pm 0.06$ | $4.0 \pm 0.1$ |
| | $k_{off}$ ($sec^{-1}$) | $(5.3 \pm 0.03) \times 10^{-4}$ | $(5.4 \pm 0.04) \times 10^{-4}$ | $(5.7 \pm 0.6) \times 10^{-5}$ | $(5.8 \pm 0.2) \times 10^{-5}$ |
| | $t_{1/2}$ (mins) | $21.4 \pm 0.07$ | $21.5 \pm 0.06$ | $205 \pm 10$ | $198 \pm 8$ |
| | $K_d$ ($\mu M$) | 0.015 | 0.045 | 28.3 | 14.6 |
| TAN | $k_2/K$ ($M^{-1}.sec^{-1}$) | $(0.9 \pm 0.02) \times 10^4$ | $(0.2 \pm 0.02) \times 10^4$ | $17 \pm 0.7$ | $15.2 \pm 0.1$ |
| | $k_{off}$ ($sec^{-1}$) | $(1.0 \pm 0.04) \times 10^{-4}$ | $(1.1 \pm 0.03) \times 10^{-4}$ | $(5.0 \pm 0.4) \times 10^{-5}$ | $(5.2 \pm 0.3) \times 10^{-5}$ |
| | $t_{1/2}$ (mins) | $104 \pm 2.3$ | $106 \pm 2.3$ | $231 \pm 19.4$ | $222.3 \pm 13.7$ |
| | $K_d$ ($\mu M$) | 0.011 | 0.055 | 2.9 | 3.4 |
| LED | $k_2/K$ ($M^{-1}.sec^{-1}$) | $^a(2.9 \pm 0.07) \times 10^4$ | $(3.1 \pm 0.8) \times 10^4$ | $(5.2 \pm 0.6) \times 10^2$ | $(4.3 \pm 0.3) \times 10^2$ |
| | $k_{off}$ ($sec^{-1}$) | $^a(2.5 \pm 0.1) \times 10^{-4}$ | $(3.7 \pm 0.1) \times 10^{-4}$ | $(6.9 \pm 0.2) \times 10^{-4}$ | $(6.3 \pm 1.0) \times 10^{-4}$ |
| | $t_{1/2}$ (mins) | $^a46 \pm 2$ | $32 \pm 1$ | $16.7 \pm 0.5$ | $18.6 \pm 2.7$ |
| | $K_d$ ($\mu M$) | 0.009 | 0.012 | 1.3 | 1.5 |

Data are reported as the mean ± standard deviation, (n = 3)
<sup>a</sup>: Kinetic data for Ledaborbactam with KPC-2<sup>wt</sup> was previously reported<sup>31</sup>.

**Table 7.** Kinetic parameters for inhibition of KPC-3 variants.

| Inhibitor | Parameter | wt | V240G | D179Y | D179Y T243M |
| --- | --- | --- | --- | --- | --- |
| AVI | $k_2/K$ ($M^{-1}.sec^{-1}$) | $(4.0 \pm 0.3) \times 10^4$ | $(2.4 \pm 0.4) \times 10^4$ | $(7.0 \pm 0.5) \times 10^3$ | $(4.6 \pm 0.8) \times 10^3$ |
| | $k_{off}$ ($sec^{-1}$) | $(1.1 \pm 0.009) \times 10^{-3}$ | $(1.1 \pm 0.009) \times 10^{-3}$ | $(9.0 \pm 0.7) \times 10^{-4}$ | $(5.2 \pm 0.7) \times 10^{-4}$ |
| | $t_{1/2}$ (mins) | $10.1 \pm 0.1$ | $10.7 \pm 0.1$ | $12.8 \pm 0.9$ | $22.5 \pm 3.5$ |
| | $K_d$ ($\mu M$) | 0.028 | 0.046 | 0.129 | 0.113 |
| TAN | $k_2/K$ ($M^{-1}.sec^{-1}$) | $(3.9 \pm 0.4) \times 10^4$ | $(4.4 \pm 0.9) \times 10^4$ | $(2.8 \pm 0.2) \times 10^3$ | $(4.4 \pm 0.7) \times 10^3$ |
| | $k_{off}$ ($sec^{-1}$) | $(3.5 \pm 0.4) \times 10^{-4}$ | $(8.0 \pm 0.4) \times 10^{-4}$ | $(8.1 \pm 0.7) \times 10^{-4}$ | $(8.5 \pm 2.5) \times 10^{-4}$ |
| | $t_{1/2}$ (mins) | $34 \pm 4$ | $14.4 \pm 0.7$ | $14.3 \pm 1.2$ | $14.6 \pm 5$ |
| | $K_d$ ($\mu M$) | 0.009 | 0.018 | 0.290 | 0.193 |
| LED | $k_2/K$ ($M^{-1}.sec^{-1}$) | $(2.7 \pm 0.05) \times 10^3$ | $(1.3 \pm 0.4) \times 10^4$ | $(5.4 \pm 2.4) \times 10^2$ | $(1.0 \pm 0.3) \times 10^3$ |
| | $k_{off}$ ($sec^{-1}$) | $(5.9 \pm 0.06) \times 10^{-4}$ | $(6.0 \pm 0.1) \times 10^{-4}$ | $(6.7 \pm 0.1) \times 10^{-4}$ | $(6.5 \pm 0.8) \times 10^{-4}$ |
| | $t_{1/2}$ (mins) | $19.6 \pm 0.2$ | $19.3 \pm 0.4$ | $17.2 \pm 0.3$ | $18.1 \pm 2.2$ |
| | $K_d$ ($\mu M$) | 0.22 | 0.046 | 1.25 | 0.62 |

Data are reported as the mean ± standard deviation, (n = 3)

In agreement with Compain and Arthur^23^, *k*^2^/*K* for AVI was reduced 18,000-fold with KPC-2^D179Y^ and 9,000-fold with KPC-2^D179Y^ ^T243M^ (Table 6). Inactivation efficiencies of KPC-2^D179Y^ and KPC-2^D179Y^ ^T243M^ by TAN were also reduced, but to a much lesser degree than AVI, roughly 500- and 600-fold, respectively. Estimations of *K*^d^ indicated higher affinity of these variants for TAN than for AVI (Table 6). Inhibition of KPC-2^D179Y^ and KPC-2^D179Y^ ^T243M^ by LED was even less impacted than TAN – *k_2_/K* was reduced ∼ 60-fold and ∼ 70-fold for KPC-2^D179Y^ and KPC-2^D179Y^ ^T243M^, respectively, which was reflected in improved *K*^d^s compared to AVI or TAN. These findings confirm that TAN and LED inhibitory activities are less affected by the D179Y substitution in KPC-2, which contributes to greater rescue of the antimicrobial activity by the partner BL compared to AVI (e.g. rescue of CAZ activity by TAN, Table 1).

TAN, LED and AVI inactivated KPC-3^D179Y^ and KPC-3^D179Y^ ^T243M^ with modestly reduced (∼2-14-fold) efficiencies relative to KPC-3^wt^ (Table 7). Similar findings for AVI have been reported by Shapiro and colleagues^37^. Dissociation kinetics for TAN, AVI and LED were faster with KPC-3^D179Y^ and KPC-3^D179Y^ ^T243M^ than for corresponding KPC-2 variants, and while AVI and TAN underwent slower dissociation from KPC-2^D179Y^ variants compared to KPC-2^wt^, off-rates for KPC-3 D179Y variants were similar to KPC-3^wt^ (Table 7). Therefore, the D179Y substitution produced different impacts on the inhibition of KPC-3 versus KPC-2. Given the moderate effect of this substitution on KPC-3 inhibition, the observed differences between CTB-LED, FEP-TAN and CAZ-AVI antibacterial activity against D179Y-containing KPC-3 variant-expressing strains appear to be primarily due to the different reactivities of these variants with CAZ versus FEP, or CTB.

### Molecular basis for the impact of the D179Y substitution on inhibition of KPC

The finding that the D179Y mutation caused different effects on the inhibitory properties of TAN, LED and AVI underscored the distinct modes of inhibitor binding to these variants. X-ray crystal structures have been solved for complexes of AVI^38^ and TAN^39^ with KPC-2. For AVI, the sulfate moiety participates in hydrogen bonding interactions with S130, T235 and T237, and the carbonyl with backbone nitrogens from S70 and T237. The amide moiety makes additional hydrogen bonds with N132, as well as with the deacylation water (positioned by E166 and N170), while making van der Waals interactions with E166 and N170. Additionally, W105, thought to be important for β-lactam substrate recognition^40^, was observed making modest van der Waals interactions with the 6-membered piperidine ring from AVI.TAN interacts with essentially the same active site residues as AVI^38,39^. The bicyclic boronate core interacts with S130, T235 and T237, and makes hydrophobic interactions with the indole of W105. The acetamido moiety bridging the cyclohexane ring to the bicyclic boronate core engages N132 and T237, and the hydroxyl group on the boron interacts with the backbone amides of T237 and S70, along with the deacylating water. The orientation of the cyclohexane ring was undefined owing to the inherent flexibility of this moiety^39^.

The recently determined crystal structure for KPC-2^D179Y^ showed little deviation in the orientation of key active site residues, except for W105 which was flipped perpendicular to its orientation in KPC-2^wt^ ^18^ (Figure. 3). Also, the absence of electron density for Ω-loop residues in the KPC-2^D179Y^ structure was consistent with a disordered Ω-loop, which has been further supported by NMR spectroscopy findings^41^. Accordingly, the loss of the inhibitor interactions dependent on this loop would likely account for the diminished inhibition of this variant.

**Figure 3.**
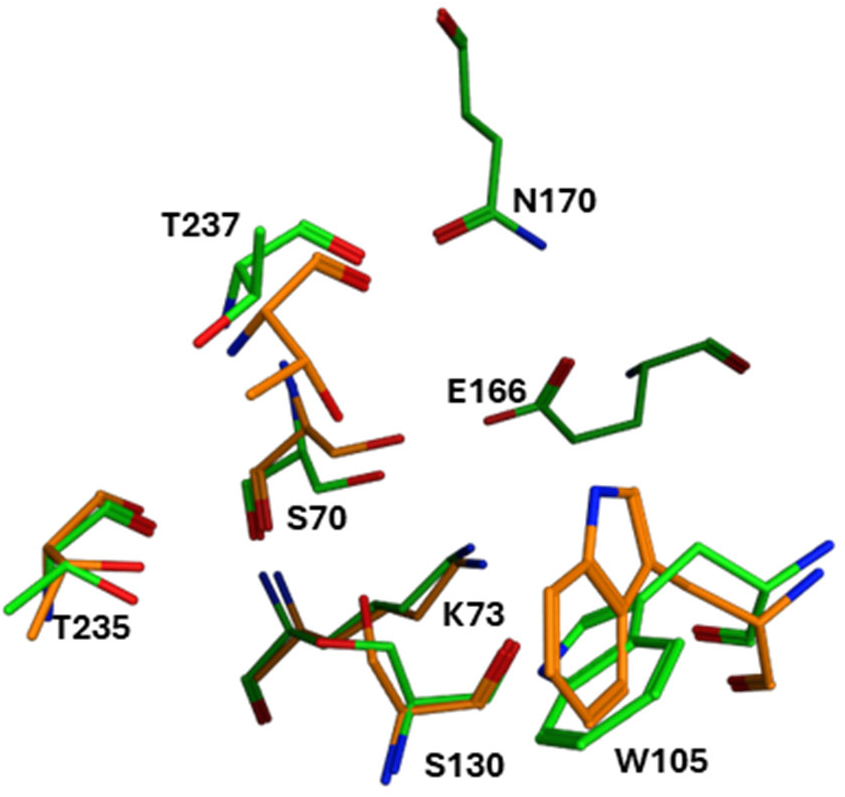
Comparison of active site residues for KPC-2^wt^ (green) and KPC-2^D179Y^ (orange).

Covalent docking of TAN to KPC-2^D179Y^ implied that critical interactions with T235 and T237 could be compromised (Figure. 4A, 4B). It has been suggested that the disorder of the Ω-omega loop in this variant would be transmitted to the region containing W105^39^, thus perturbing inhibitor interactions with this residue.

**Figure 4.**
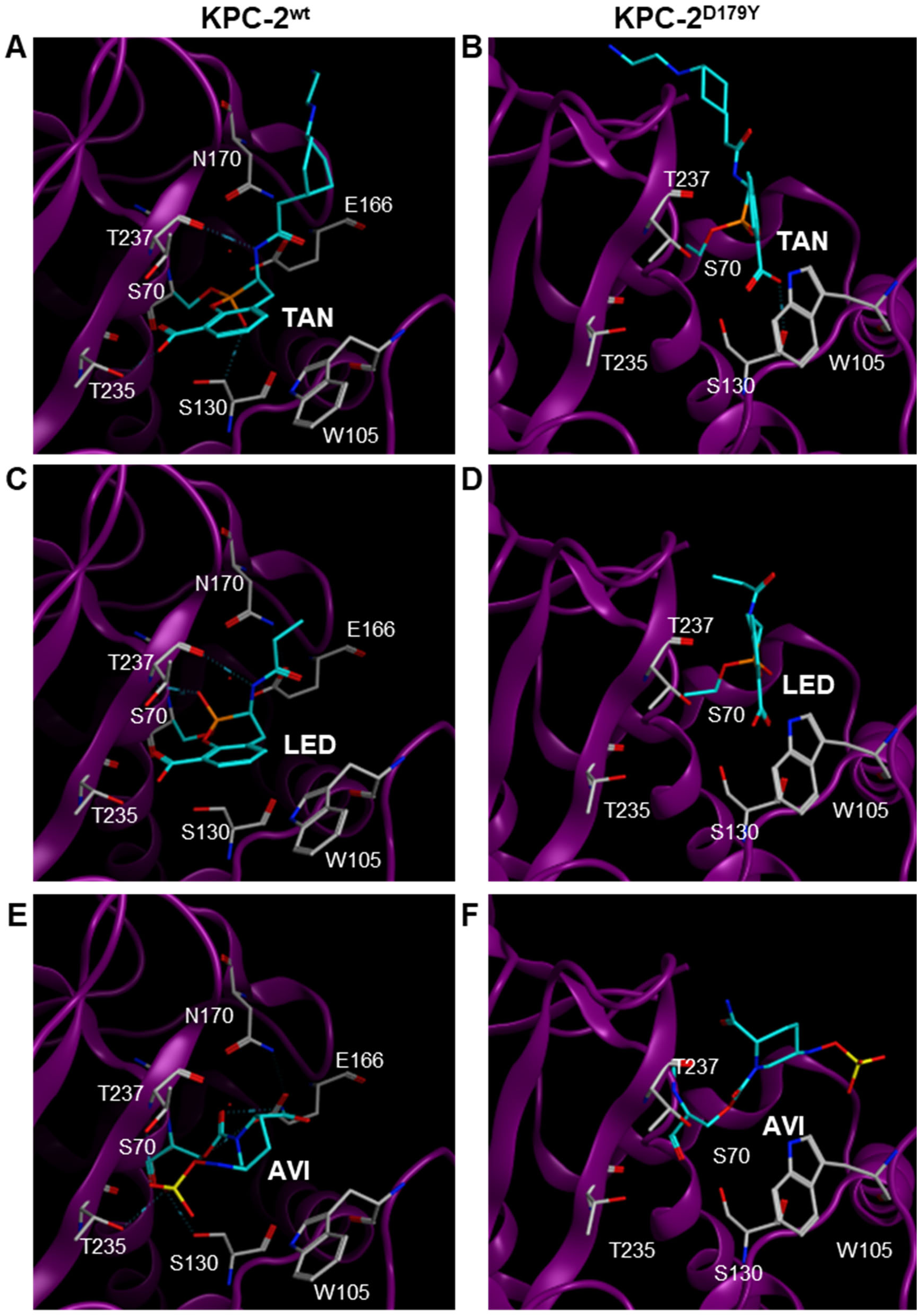
Molecular docking of TAN, LED and AVI with KPC-2^wt^ (A, C and E) and KPC-2^D179Y^ (B, D and F).

Accordingly, the bicyclic core of TAN appeared displaced from its interaction with W105 (Fig. 4B). For AVI, interactions with W105 in KPC-2^D179Y^ are completely lost, along with the interaction with S130 (Fig. 4E, 4F). Modelling of LED in the KPC-2^wt^ active site showed its bicyclic core engaged in similar interactions to those observed with TAN, and important interactions with T235, T237 and especially W105 in KPC-2^D179Y^ were disrupted in an analogous manner (Fig. 4C, 4D).

## Discussion

Infections by Gram-negative bacteria expressing carbapenemases pose a serious threat to global health for which treatment options have been limited to less desirable agents that feature unfavourable toxicity or efficacy profiles. Chief among such carbapenemases are KPCs, but the combination of CAZ with AVI has provided an effective, well-tolerated therapy for infections by KPC-producing bacteria. Unfortunately, KPC variant-mediated CAZ-AVI resistance has quickly evolved in the clinic, complicating empiric and follow-up therapies^42^. *Bla*^KPC^ mutations causing amino acid changes in the Ω-loop, especially at D179, were among the first and most commonly identified, and additional variants are being identified with more frequency^21^. In this study, we showed that bicyclic boronate BLIs TAN and LED restored cephalosporin activity against engineered CAZ-AVI-resistant

*E. coli* strains expressing KPC-variants by circumventing distinct effects on the hydrolytic activity and inhibition of KPC-2 versus KPC-3. A key finding from this work was that the hydrolysis and interactions of CAZ were more profoundly affected than FEP or CTB. Of the three variants, KPC-2/ KPC-3 possessing the V240G substitution caused the least resistance to CAZ-AVI. It was apparent that the highest concentrations of CAZ, FEP or CTB, used in MICs assays (256 µg/mL) would result in intra-cellular concentrations well below the respective *K*^M^s for V240G-containing variants, or corresponding wild-type enzymes (see Tables 3 & 4). Hence, these enzymes would be operating below maximal efficiency, which would enable BLI rescue in cells. Even so, it was notable that FEP or CTB activity was more effectively rescued than CAZ in a V240G variant expression background, which likely reflects higher *k*^cat^ for CAZ by these variants relative to wild-type enzymes.

Effects from substitutions at D179 on CAZ hydrolysis by KPC, and on the antibacterial activity of CAZ or CAZ-AVI, have been studied previously^18,24,26,37^. This residue is engaged in a salt bridge with R164 that stabilizes the Ω-loop, and substitutions at D179 induce disorder in the loop region that is thought to allow for better accommodation of CAZ in the enzyme active site while facilitating its prolonged interaction with the enzyme^24^. This aligns with the reductions in *K*^M^ (and presumably *k*^cat^) observed here for the D179Y-containing variants with CAZ, as well as with FEP, and is consistent with impaired deacylation that leads to the prolonged existence of the acyl-enzyme complex^24,43^. Therefore, the increased CAZ affinity for D179Y-containing variants relative to the wild-type enzyme affirms the proposed role of enhanced substrate affinity in CAZ-AVI resistance mediated by KPC^D179Y^ or KPC^D179Y^ ^T243M^. Importantly, FEP was found to have significantly weaker affinity relative to CAZ for D179Y-containing variants, while CTB lacked measurable affinity by the approaches used here. This highlights the influence of BL molecular structure, particularly the presence and nature of the R2 group (Fig. 5), on binding to KPC variants. Thus, poor binding affinity and/ or weak hydrolysis by D179Y-containing variants enables virtually complete rescue of FEP or CTB antibacterial activity by TAN or LED in expressing strains.

**Figure 5.**
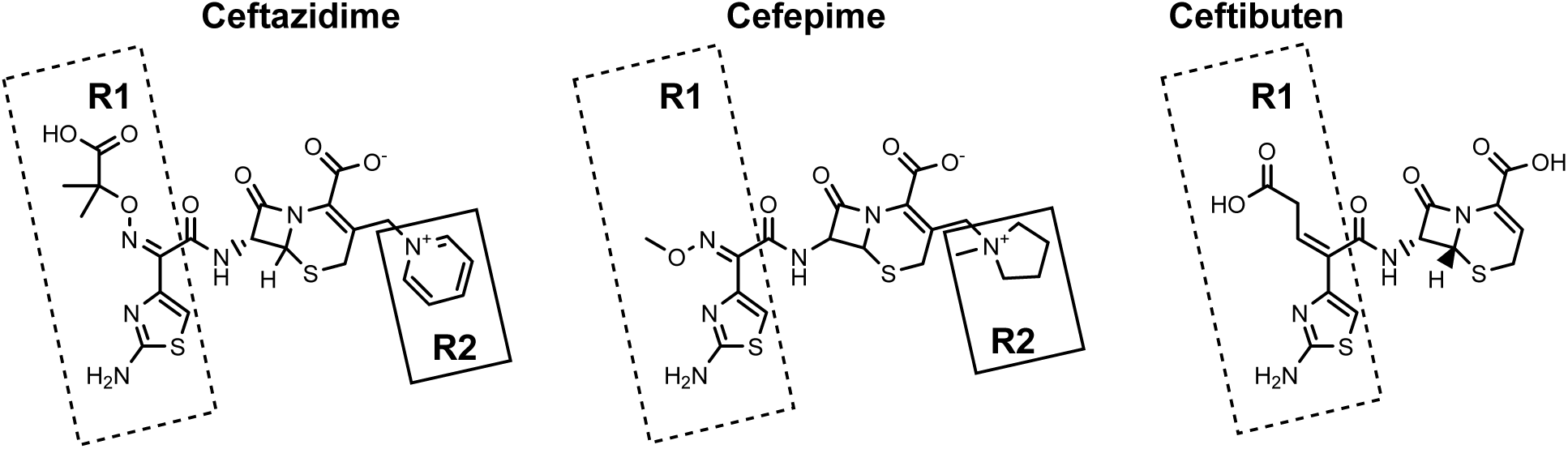
Structures of the cephalosporins used in this study, with R1 (dashed lines) and R2 (solid lines) groups indicated.

Molecular docking experiments indicated possible alterations in inhibitor binding to KPC-2^D179Y^ that could account for the diminished inhibition of variants with this amino acid substitution. W105 contributes important hydrophobic interactions to inhibitor binding in the wild-type enzyme, and the loss of AVI interactions with this residue provides a plausible explanation for the reduced efficiency with which the BLIs inactivate KPC-2^D179Y^, as well as the higher inactivation efficiency of TAN and LED compared to AVI. Like TAN, LED took longer to form a covalent adduct with KPC-2^D179Y^ than with the wild-type enzyme, but KPC-2^D179Y^ was more efficiently inactivated by LED than by TAN. A possible explanation could be that the propionamide moiety in LED experiences less steric hindrance from Y179 compared to the cyclohexyl diamine in TAN (or the piperidine ring in AVI). Indeed, the inherent flexibility of the cyclohexyl diamine in TAN may hinder adduct formation to KPC-2^D197Y^. This could be exacerbated by disorder in the Ω-loop in KPC^D179Y^ that may encourage unfavourable steric clashes with the cyclohexyl diamine in TAN (or the piperidine ring in AVI).

LED inactivated KPC-2^D179Y^ and KPC-2^D179Y^ ^T243M^ with greater efficiencies than AVI or TAN, but inactivated KPC-3 D179Y-containing variants with lower efficiencies than AVI or TAN. Moreover, AVI, TAN and LED all retained significant inhibition of KPC-3 variants. We have shown here that KPC-2 and KPC-3 variants exhibited different catalytic properties with respect to BL hydrolysis, but it is not immediately obvious why the D179Y substitution produces distinct effects on the hydrolytic activity or inhibition of KPC-2 versus KPC-3. KPC-2 and KPC-3 differ at a single position – residue 274 is a histidine in KPC-2 and a tyrosine in KPC-3. It has been suggested that Y274 in KPC-3 supplies additional binding interactions to CAZ^34^, which can be envisioned to apply to inhibitor binding. Alternatively, this residue could modulate protein dynamics with implications for substate/ inhibitor binding. Additional biochemical, structural and biophysical studies are needed to elucidate the details underlying this difference.

## Summary and conclusions

FEP-TAN and CTB-LED are potently active against *E. coli* strains expressing KPC variants that confer resistance to CAZ-AVI. The biochemical basis for this activity was examined, revealing effects on hydrolytic activity and inhibition that account for the observed BL or BL/BLI susceptibilities against KPC-2 versus KPC-3 variant-expressing strains. D179Y-containing variants demonstrated an increased affinity for CAZ or FEP, especially the former, and CTB, like FEP, was a poorer substrate than CAZ and showed weak affinity for these variants. In KPC-2, the D179Y substitution led to significantly diminished inhibition by AVI, TAN and LED, with AVI being most impaired, whereas inhibition of corresponding KPC-3 variants was modestly impacted. Therefore, FEP-TAN and CTB-LED are active against strains producing KPC-2 D179Y variants due to sufficient inhibition by the boronate inhibitors along with weak hydrolysis and poorer affinity of the partner cephalosporins. FEP-TAN and CTB-LED activity against D179Y-containing KPC-3 variant-producing strains, however, is due primarily to the weaker affinity of these enzymes for FEP or CTB compared to CAZ, and poor hydrolysis in the case of CTB. Overall, the findings show that strains expressing KPC variants of clinical importance are within the spectrum of antibacterial activity for the cephalosporin-bicyclic boronate BLI combinations and offer evidence supporting their use for treating infections caused by KPC-producing Enterobacterales.

## Materials and Methods

### Bacterial strains and microbiological assays

Isogenic *E. coli* strains engineered for periplasmic expression of KPC-2, and KPC-3 variants were constructed as previously described^28^. MICs were determined by broth microdilution according to CLSI standard methods^44,45^.

### Protein expression and purification

DNA sequences encoding full-length KPC-2, and variants were cloned into pET9A plasmid and plasmid constructs were used to transform *E. coli* BL21 (DE3) cells. Cells were cultured in LB medium supplemented with 35 µg/ml kanamycin to an OD^600^ of 0.6 – 0.8, after which cultures were cooled, then protein expression induced with 1 mM isopropyl-β-D-1-thiogalactopyranoside (IPTG) for 16 hours at 18°C. Cells were harvested by centrifugation at 6,000 × g and the periplasmic contents, containing mature KPC-2 proteins, were extracted by cold osmotic shock^46^ in the presence of 0.02 mg/mL lysozyme. KPC-2 proteins were purified using ion-exchange chromatography, followed by size-exclusion chromatography, as previously described^28^.

The sequence encoding the mature, periplasmic domain of KPC-3 and variants (residues 27 – 293) were cloned into pET28a plasmid and the resulting constructs transformed into *E. coli* BL21 (DE3) cells. Cells were grown at 37°C in 1L of LB medium supplemented with 35 µg/ml kanamycin to an OD^600^ of 0.6 – 0.8, after which cultures were cooled, then protein expression was induced with 1 mM IPTG for 16 hours at 18°C. Cells were harvested by centrifugation (6,000 × g), resuspended in buffer A (20 mM Tris (pH 7.5), 300 mM NaCl, 5% glycerol) supplemented with EDTA-free protease inhibitor, then lysed by sonication (Branson Sonifier 250). Lysates were clarified by centrifugation at 14,000 × g, and KPC-3 proteins purified using immobilized metal ion chromatography, by applying lysates to 5 ml of Ni-NTA resin (ThermoFisher) equilibrated with buffer A. The resin was washed with 10 mM imidazole in buffer A, then His-tagged protein was eluted in a stepwise fashion with increasing concentrations of imidazole (50 mM, 75 mM, 250 mM, 500 mM) in buffer A. Fractions containing pure protein were pooled, then concentrated and the buffer exchanged for 20 mM Tris (pH 7.5), 150 mM NaCl, 5% glycerol using Amicon Ultra-15 centrifugal filters before storage at -80°C.

### Enzyme assays

Measurements to determine kinetic parameters for hydrolysis, and kinetic parameters for inhibition were performed at ambient temperature in PBS (pH 7.4) containing 0.1 mg/mL bovine serum albumin, using a PowerWaveXS plate reader (BioTek).

Assays to determine Michaelis-Menten kinetic parameters were performed by mixing variable amounts of enzymes (depending on the enzyme-substrate pair), with serial dilutions of different BL substrates and continuously monitoring the reduction in absorbance associated with BL hydrolysis^33,43,47,48^. Kinetic parameters were determined by plotting initial rates of hydrolysis (v) as a function of substrate concentration [S] and fitting the data to equation 1 using GraphPad Prism.

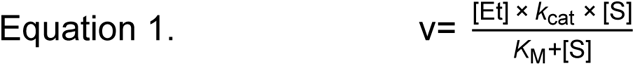

where, [Et] the enzyme concentration, *k*^cat^ the turnover number and *K*^M^ the Michaelis constant. For instances where substrate hydrolysis displayed linear kinetics (i.e. initial velocity of hydrolysis could not be saturated at obtainable substrate concentrations), *k*^cat^/*K*^M^ was estimated by fitting plots of velocity versus substrate concentration to equation 2, assuming [S] << *K* ^34^.

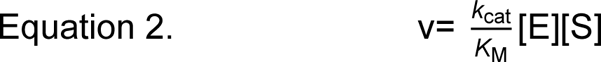

Second-order rate constants (*k*^2^/*K*) denoting the inactivation efficiency of BLIs were determined by continuously measuring CENTA (100 μM) hydrolysis^33^ in the presence of various concentrations of BLI. Progress curves were fit to equation 3 using GraphPad Prism, where A^i^ and A^0^ are the measured and basal absorbance, v^i^ and v^s^ are initial and final velocity, and t is time in seconds. *K*^2^/*K* was derived with equation 4. *K*^off^ was determined by the jump dilution method^28^. *K*^d^ was estimated from the ratio between *k*^off^ and *k*^2^/*K*.

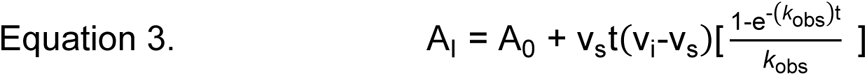

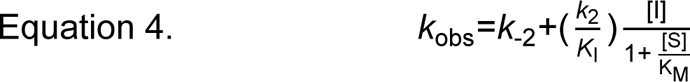

Affinities of CAZ, FEP and CTB for KPC variants were determined by competition assay^49^, with CENTA as the reporter substrate. Enzyme was added to mixtures containing CENTA and various concentrations of CAZ, FEP or CTB, and the absorbance at 405 nm was continuously monitored. The inverse of the initial velocity was plotted against the corresponding inhibitor concentrations and the data fit to a linear equation to derive *K*^i app observed^ by dividing the y-intercept by the slope of the line. *K*^i app observed^ was corrected for substrate concentration and affinity using equation 5.

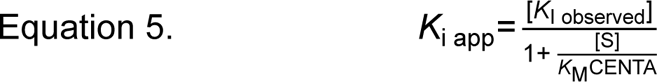

### Molecular Modelling

Complexes between KPC-2^wt^ and KPC-2^D179Y^ with AVI, TAN or LED were generated using MOE software^50^. Structures for KPC-2^wt^ (PDB: 5UL8^32^) and KPC-2^D179Y^ (PDB: 7TBX^18^) were imported into the software and prepared for docking using MOE QuickPrep. Molecular structures for AVI, TAN and LED were created in the MOE environment and energy minimized before covalent docking at the catalytic serine. Complexes with KPC-2^wt^ were validated by comparison with published co-structures for KPC-2-AVI^38^ or KPC-2-TAN^39^. For the KPC-2^D179Y^ complexes, models with the lowest binding energies are shown.

